# Type I interferon alters invasive extravillous trophoblast function

**DOI:** 10.1101/2024.03.11.584521

**Authors:** Michael K. Simoni, Seble G. Negatu, Ju Young Park, Sneha Mani, Montserrat C. Arreguin, Kevin Amses, Dan Dongeun Huh, Monica Mainigi, Kellie A. Jurado

## Abstract

Inappropriate type I interferon (IFN) signaling during embryo implantation and placentation is linked to poor pregnancy outcomes. Here, we evaluated the consequence of elevated type I IFN exposure on implantation using a biomimetic model of human implantation in an organ-on-a-chip device. We found that type I IFN reduced extravillous trophoblast (EVT) invasion capacity. Analyzing single-cell transcriptomes, we uncovered that IFN truncated endovascular EVT emergence in the implantation-on-a-chip device by stunting EVT epithelial-to-mesenchymal transition. Disruptions to the epithelial-to-mesenchymal transition is associated with the pathogenesis of preeclampsia, a life-threatening hypertensive disorder of pregnancy. Strikingly, unwarranted IFN stimulation induced genes associated with increased preeclampsia risk and a preeclamptic gene-like signature in EVTs. These dysregulated EVT phenotypes ultimately reduced EVT-mediated endothelial cell vascular remodeling in the implantation-on-a-chip device. Overall, our work indicates IFN signaling can alter EVT epithelial-to-mesenchymal transition progression which results in diminished EVT-mediated spiral artery remodeling and a preeclampsia gene signature upon sustained stimulation. Our work implicates unwarranted type I IFN as a maternal disturbance that can result in abnormal EVT function that could trigger preeclampsia.

## INTRODUCTION

Balanced immune-mediated intercellular communication underlies successful reproduction^1^. Embryo implantation into the maternal endometrium, and subsequent placentation, is a complex process where fetal trophoblast cells invade and restructure the uterine vasculature. Endometrial transformation enables low resistance blood flow to the fetus and requires a precise degree of trophoblast invasion in order to prevent placental dysfunction. Either inadequate or excessive trophoblast invasion can lead to a wide spectrum of pregnancy complications, and thereby requires fine-tuned control. Extravillous trophoblasts (EVT) arise from epithelial cytotrophoblasts with specialized migratory and invasive properties and mediate endometrial invasion and spiral artery remodeling. EVT functionality is tightly regulated by many different cell types including uterine immune cells via cytokine mediators^1,2^.

Maternal immune dysregulation and inflammation are linked to various pregnancy complications. Chronic inflammatory conditions, such as systemic lupus erythematosus (SLE), are associated with elevated risk of pregnancy loss^3,4^. Specifically, patients with lupus flares during early pregnancy are at greatest risk for pregnancy complications, with a 30-40% chance of suffering a pregnancy loss^5^. Studies have specifically linked inappropriate type I interferon (IFN) signaling during early pregnancy, when embryo implantation and placentation occur, with poor pregnancy outcomes^3,6^. Type I IFN signaling typically drives an antiviral state but can also be elevated in certain states of immune dysregulation and/or autoimmune conditions, including SLE^4^. Preeclampsia (PE) complicates 13-35% of SLE pregnancies, compared to less than 1-6% in otherwise healthy women^7^. PE is a hypertensive disorder in pregnancy that is characterized by shallow trophoblast invasion and inadequate spiral artery remodeling. Given that type I IFN is elevated in SLE patients who demonstrated pregnancy complications associated with poor trophoblast invasion, we hypothesized that type I IFN could impact EVT function.

This led us to investigate the impact of elevated type I IFN stimulation on EVT invasion and spiral artery remodeling. Using a biomimetic model of human implantation in a microengineered organ-on-a-chip device^8^, we found that exposure to type I IFN abrogated EVT invasion. Analyzing single-cell transcriptomic data obtained from cells extracted from devices, we found that unwarranted type I IFN exposure limits endovascular EVT emergence, disrupts EVT epithelial-to-mesenchymal progression and induces a preeclamptic gene signature in EVTs. These dysregulated phenotypes ultimately reduced EVT-mediated vascular remodeling in our implantation-on-a-chip device. Our findings implicate unwarranted type I IFN as a maternal disturbance that can trigger PE.

## RESULTS

### Elevated type I interferon exposure abrogates EVT invasion

To evaluate the impact of elevated type I IFN on implantation, we employed an implantation-on-a-chip (IOC) device that models the invasion of fetal extravillous trophoblasts (EVTs) into the maternal uterus^8^. The IOC device reconstructs the three-dimensional organization of the maternal-fetal interface where EVTs exhibit directional migration through an extracellular matrix (ECM) toward endometrial endothelial cells (ECs) (**Figure 1a-b**). EVT migration within the IOC device is dependent on intercellular communication with maternal endometrial ECs, highlighting its physiological relevance^8^. To begin, we seeded the IOC device with primary human first trimester EVTs isolated from clinical specimens and primary human endometrial ECs. Two days after seeding, we began daily treatments with IFN-β (1000 IU/mL), a hallmark type I IFN, by supplementing the daily media exchange of the IOC device. We next assessed the properties of EVT invasion into the ECM toward the endothelial chamber for five days (**Figure 1c**). Within 24 hours of IFN-β treatment we observed a significant reduction in EVT migration out of the trophoblast chamber and into the ECM as compared with the control (**Figure 1d-e**). This blunting of EVT invasion with IFN-β exposure persisted at day 3 and 5 (**Figure 1d-e**). At day 5 post-IFN-β exposure, we quantified the depth of invasion and the number of EVTs invaded into the ECM. Using these measurements, we observed that both the average number of invaded EVTs and the average depth of EVTs in the ECM were significantly reduced after IFN-β exposure (**Figure 1f-g**). Using a lower dose of IFN-β (100 IU/mL), we observed similar abrogation of EVT invasion with similar reductions in EVT invasion over time (**Supplemental Figure 1a & 1b**).

**FIGURE 1:**
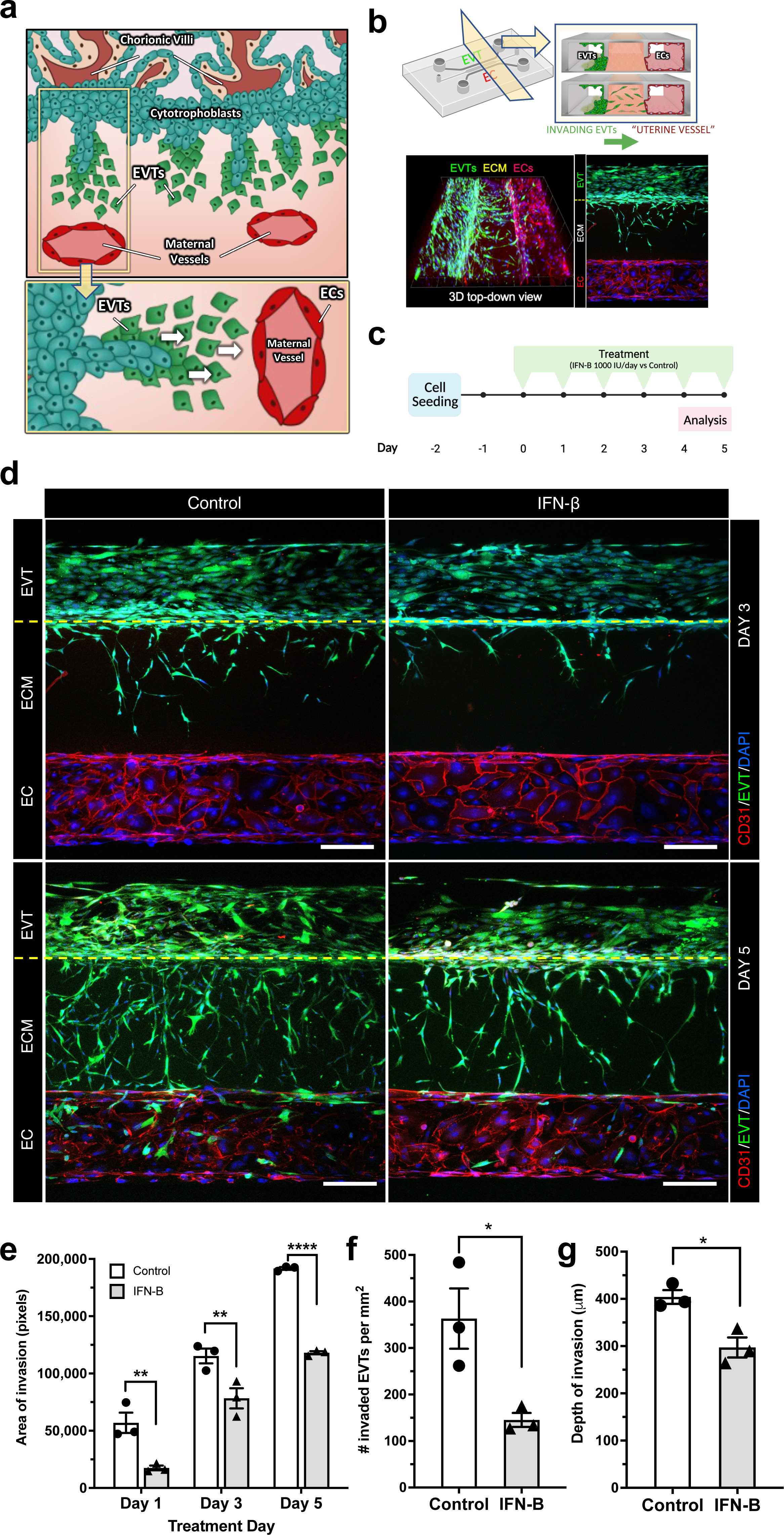
Type I interferon exposure abrogates EVT invasion. **a.** A visual portrayal of human embryo implantation, where cytotrophoblasts, present within the fetal chorionic villi, differentiate into extravillous trophoblasts (EVTs) that invade into the maternal uterus towards maternal vasculature. **b** Top left - Schematic of the implantation-on-a-chip (IOC) microfluidics device. The center and two side lanes have dimensions of 0.5 mm (width) × 0.3 mm (height) and 0.25 mm (width) × 0.3 mm (height), respectively. Top right - Compartmentalized design of the IOC device allows EVTs to migrate through an extracellular matrix (ECM) towards maternal endothelial cells (ECs). Bottom row - 3D (left) and 2D top-down (right) representative images of EVTs (green - CellTracker Green) migrating across the ECM hydrogel towards maternal ECs (red – CD31). **c.** Graphic of experimental timeline shown in days with start of type I interferon (IFN-β) treatment at Day 0. **d.** Representative image of IOC device at day 3 (top row) and day 5 (bottom row) with EVTs (green - CellTracker Green) migrating towards maternal ECs (red – CD31) in control and after IFN-β (1000 IU/mL) exposure; scale bars, 200 μm. Representative images are from three independent experiments. **e.** Quantification of EVT area invasion at Day 1, 3 and 5. **f-g.** Quantification of number of invaded EVTs (**f.**) and depth of invasion (**g.**). Data are presented as mean ± SEM. One-way ANOVA with Tukey’s multiple comparison test. **\*** = p < 0.05; **\*\*** = p < 0.01; **\*\*\*\*** = p < 0.0001 (*n* = 3 independent devices per group).

Next, we evaluated if IFN-β exposure impacted cellular proliferation. Previously, we had found that primary EVTs proliferate on the IOC device in the presence of endometrial endothelial cells^8^. Here, we noted a decrease in Ki-67 expression at 3- and 5-days post IFN-β exposure as compared with control (**Supplemental Figure 1c**), indicating a reduction in EVT proliferative capacity after prolonged IFN-β exposure. Taken together, these findings indicate that elevated type I IFN exposure impairs EVT invasive capacity.

### Elevated type I interferon limits endovascular extravillous trophoblast emergence

To investigate how elevated IFN-β exposure is impacting EVT invasion, we next determined the transcriptomes of cells isolated from the IOC device via scRNA sequencing. At 5-days post-IFN- β exposure or PBS control, we extracted 38,495 cells from all layers of thirty-six IOC devices and processed for 10X Genomics. Raw counts were filtered for empty cells, doublets, and other quality control parameters. We used Seurat (v4.0.4) to combine all samples into a single object, and as expected UMAP dimensionality reduction yielded two primary groups of cells (**Figure 2a**). Cellular markers unique to EVTs and ECs were used to distinguish the identity of each primary cell group. EVT clusters were distinguished by expression of trophoblast glycoprotein (*TPBG*) and placenta-specific protein 9 (*PLAC9*) while EC clusters were determined by CD31 (*PECAM1*) and Von Willebrand Factor (*VWF*) expression (**Figure 2b**). Clustertree analysis with a resolution of 0.2, resulted in four clusters within the EVT group, and two within the EC group, indicating cell state heterogeneity amongst EVTs and ECs.

**FIGURE 2:**
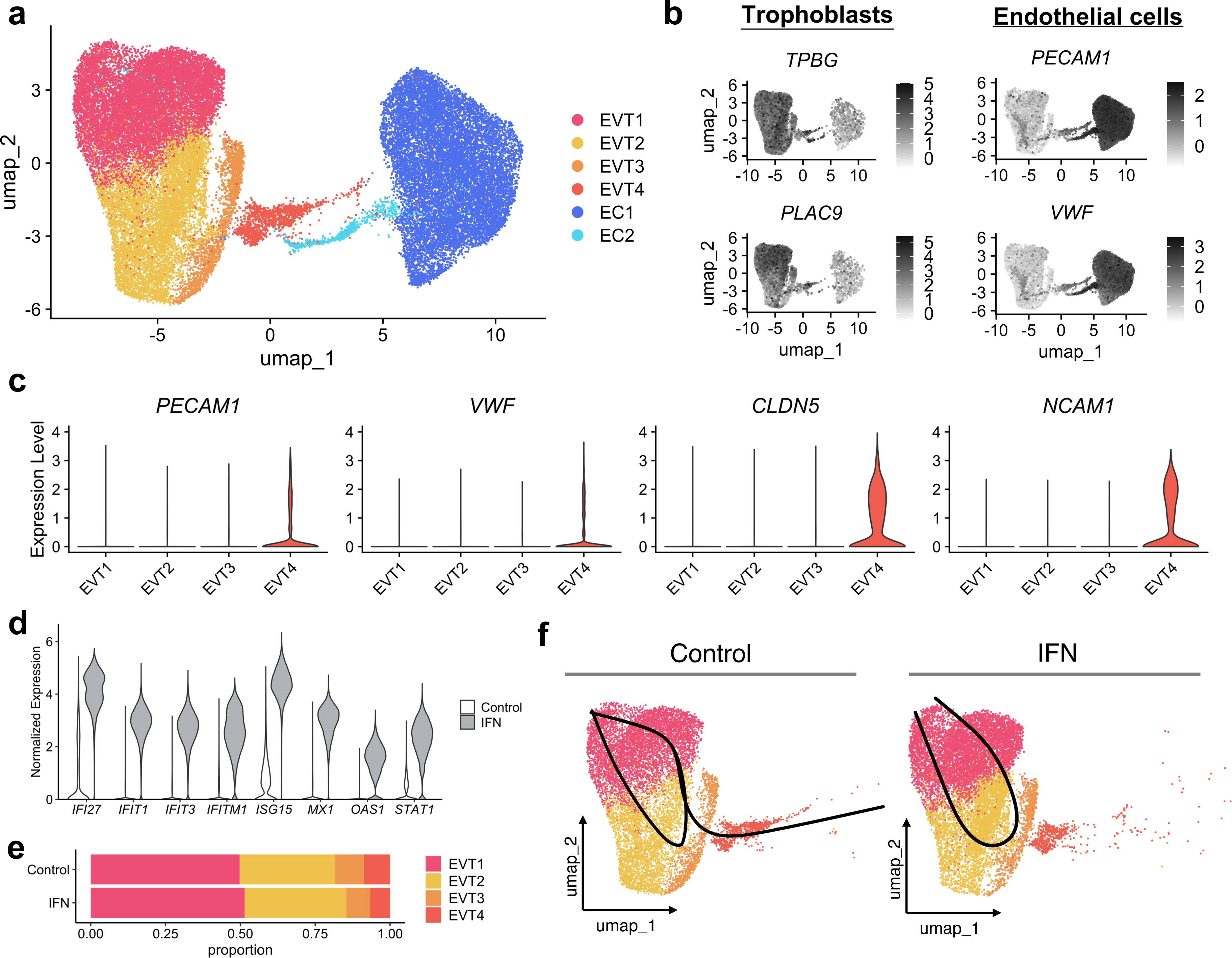
Elevated type I interferon limits endovascular extravillous trophoblast emergence. **a.** Uniform manifold approximation and projection (UMAP) plot of scRNA-seq of cells within 36 implantation-on-chip devices (*n* = 38,495 cells) colored by cell type and state. **b.** UMAP plot of marker genes characteristic of trophoblasts (left) and endothelial cells (right). **c.** Violin plots showing normalized and log-transformed expression of genes characteristic of endovascular EVTs in individual EVT subsets (x-axis). **d.** Violin plots showing normalized expression of representative interferon stimulated genes (*x*-axis) from scRNA-seq data in all cells in the IOC device separated by control (white) and type I interferon (IFN) exposed cells (grey). **e.** Bar plot representing the proportion (%) of EVT subsets separated by control and IFN stimulated cells. **f.** Minimum spanning tree computed by Slingshot, visualized on the UMAP of EVT subsets; separated by control and type I IFN stimulated cells.

Upon trophoblast invasion into the maternal endometrium, EVTs remodel and replace the maternal endothelium of uterine arteries in order to promote low-resistance blood flow to the placenta. EVTs will adopt an endothelial-like state during endovascular invasion^9,10^. Fascinatingly, the UMAP yielded a convergence of transcriptomes between the two primary cell types (**Figure 2a**). Furthermore, the cellular cluster at convergence, EVT4, uniquely expressed endothelial-smarkers *PECAM1, VWF, and CLDN5* along with the well-described marker of endovascular trophoblasts, *NCAM1* (**Figure 2c**)^9,11^. To further benchmark EVT4s, we integrated snRNA-seq data from ∼2000 interstitial and endovascular EVTs from a reference human implantation site^12^, and identified substantial overlap with EVT4s (cluster 3 in **Supplemental Figure 2**). Incredibly, these data support that endovascular EVTs emerge within the IOC device.

When IFN-β binds to the type I IFN receptor complex (IFNAR), it initiates signaling through the Janus kinase-signal transducer and activator of transcription (JAK-STAT) pathway and leads to the transcription of many IFN-stimulated genes (ISGs). Indeed, we observed expression of several canonical ISGs (*IFI27, IFIT1, IFIT3, IFITM1, ISG15, MX1, OAS1, STAT1*) across all EVT and EC clusters that were exposed to IFN-β as compared to control (**Figure 2d**), indicating a robust response to the cytokine stimulation. We next quantified the proportion of cells within clusters to assess the impact of type I IFN exposure on cell type heterogeneity. Consistent with invasion assays (**Figure 1**), the changes were primarily observed amongst the EVT clusters that represent the convergence between the two cell types: EVT3 and EVT4 (**Figure 2a**). EVT4 cluster exhibited the largest proportional difference of all EVT cells, decreasing by 22% after IFN exposure, followed by EVT3, which decreased by 17% (**Figure 2e**). We next used Slingshot to resolve subset emergence in pseudotime within both control EVTs and in IFN-exposed EVTs (**Figure 2f**)^13^. Intriguingly, trajectory analysis predicted a loss of trajectory toward EVT4 in the type I IFN treated EVTs compared to control EVTs (**Figure 2f**). This truncation in emergence progression was also supported by an alternative trajectory analysis platform, Monocle3 (**Supplemental Figure 3**)^14^. Collectively, from these data we conclude that elevated type I IFN exposure limited the emergence of endovascular EVTs in the IOC device.

### Elevated type I interferon alters epithelial-to-mesenchymal transition and promotes a preeclamptic gene signature in extravillous trophoblasts

Epithelial-to-mesenchymal transition (EMT) is a dynamic transformation of cellular organization from an epithelial state to a mesenchymal phenotype and leads to the functional acquisition of cellular migration and invasion. Cytotrophoblast to EVT differentiation is a unique form of EMT where EVTs exist along a spectrum of mixed epithelial and mesenchymal states that can readily interconvert (**Figure 3a**)^15,16^. EMT is dynamic throughout gestation where first trimester EVTs display more developed mesenchymal features during active endometrial remodeling as compared with third trimester EVTs which comparatively present with more epithelial features^16^. This epithelial-to-mesenchymal plasticity ensures that EVT migration and invasion can be regulated by decidual checkpoints to avoid pathogenic outcomes6.

**FIGURE 3:**
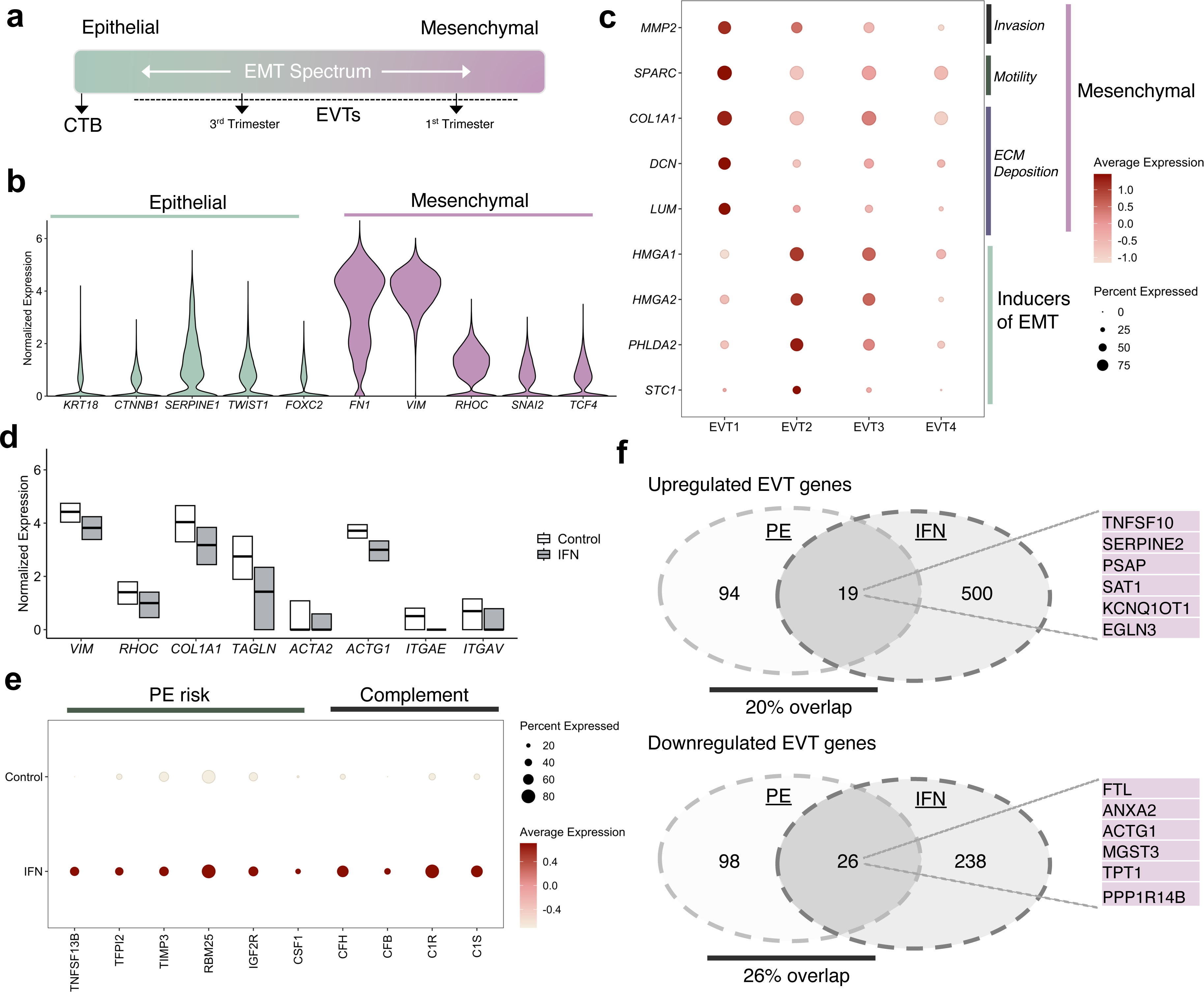
Elevated type I interferon alters epithelial-to-mesenchymal transition and promotes a preeclamptic gene signature in extravillous trophoblasts. **a.** Schematic describing the dynamic spectrum of epithelial-to-mesenchymal transition (EMT) in cytotrophoblast to extravillous trophoblasts (EVTs) transition. Small arrows indicate first trimester EVTs display more developed mesenchymal features during active endometrial remodeling as compared with third trimester EVTs **b.** Violin plots showing normalized expression of representative canonical epithelial genes (left) and mesenchymal genes (right) in all EVT cells. **c.** Dot plots showing normalized, log-transformed, and variance-scaled expression of genes characteristic of mesenchymal genes and regulatory, inducers of EMT (y-axis) in individual EVT subsets (x-axis). **d.** Box plot showing normalized of classical mesenchymal genes in all EVT cells separated by control (white) and IFN stimulated cells (grey). **e.** Dot plots showing normalized, log-transformed, and variance-scaled expression of preeclampsia risk factors (x-axis) in all EVT separated by control and IFN stimulated cells (y-axis). **f.** Venn diagram of dysregulated genes in EVTs from PE patients with early- and late-onset preeclampsia with our list of differentially expressed genes in IFN-exposed EVTs (upregulated and downregulated genes had a false detection rate less 5% and log fold change greater than one). Representative genes that overlap are listed to the right.

All EVT clusters expressed high levels of mesenchymal genes (*FN1, VIM, RHOC*) and transcription factors (*SNAI2, TCF-4*) in addition to lower levels of epithelial genes (*KRT18, CTNNB1, SERPINE1)* and transcription factors (*TWIST, FOXC2*) (**Figure 3b**). We noted that EVT1s expressed higher levels of genes characteristic of a more developed mesenchymal phenotype, including genes involved in invasion (*MMP2*) and motility (*SPARC*), extracellular matrix deposition (*DCN, LUM*, *COL1A1*) (**Figure 3c**). EVT2 more distinctly expressed regulatory, inducers of EMT (*HMGA1, HMGA2*, *PHLDA2*, *STC1*)^17–20^. Interestingly, the trajectory analysis on EVTs in **Figure 2f** proposes a pathway of interconversion between EVT1 and EVT2 clusters and thereby might represent the plasticity of EVTs along the epithelial-to-mesenchymal spectrum (**Figure 3a**).

Notably, exposure to type I IFN resulted in several classical mesenchymal genes to be considerably reduced in all EVTs (**Figure 3d**). This included decreased expression of canonical mesenchymal markers such as *VIM* and *RHOC* and extracellular matrix proteins (*COL1A1*). Furthermore, we saw reduced expression of genes involved in cell motility (TAGLN, ACTA2, *ACTG1*), and mesenchymal integrins (*ITGAE, ITGAV*). From this, we deduce that unwarranted type I IFN leads to a stunted epithelial-to-mesenchymal progression in EVTs.

Disruptions to epithelial-to-mesenchymal transition is associated with the pathogenesis of PE^21,22^. PE is characterized by shallow trophoblast invasion and a failure to adequately remodel spiral arteries, therefore we next evaluated genes known to be associated with PE pathophysiology. We found heightened expression of several genes associated with increased PE risk including *TNFSF13B, TFPI2, TIMP3, RBM25, IGF2R and CSF1*^21,23–26^ in IFN-EVTs compared with control EVTs (**Figure 3e**). We also observed increased expression of complement activation components that have specifically been implicated in PE pathogenesis (*CFH, CFB, C1R, C1S*) in IFN-exposed EVTs^27,28^. Given these associations, we next compared our list of differentially expressed genes in IFN-exposed EVTs with a list of dysregulated genes in EVTs from single-cell transcriptomes of EVTs in PE patients recently published by Admati et al.^29^. We identified noteworthy overlap with 20% of upregulated genes and 26% of downregulated genes shared with dysregulated EVT genes found in PE patients (**Figure 3f**). Taken together, these data indicate elevated type I IFN exposure disrupts EVT-EMT progression and result in a PE-like gene signature in EVTs.

### Elevated type I interferon limits EVT-mediated vascular remodeling in IOC device

Spiral artery remodeling is critical for enabling adequate blood supply to the fetus and involves the replacement of luminal ECs in uterine arteries with endovascular trophoblast. Physiological transformation of maternal vasculature is a dynamic process that is mediated by EVTs. Vascular disruption and endothelial apoptosis are two major features of spiral artery remodeling upon endovascular EVT invasion. Notably, the IOC device models these processes in a physiologically relevant manner^8^. Since we observed that type I IFN exposure impaired EVT invasion in the IOC and reduced endovascular EVT emergence and overall EVT quality, we next aimed to ask if this ultimately impacted EVT-induced vascular remodeling. To begin, we resolved the most likely trajectory of cell state emergence in pseudotime amongst all EC cells via Slingshot. We found the primary cluster of EC1 transitions into the minor EC2 cluster, indicating EC2s are undergoing a transformation of cell state (**Figure 4a**). In order to ask if EC2s are representative of ECs undergoing remodeling, we next assessed expression of an important adhesion protein commonly used to distinguish vascular disruption during implantation, vascular endothelial (VE-)cadherin (*CDH5)*. We found that EC2 had decreased *CDH5* levels as compared with EC1 (**Figure 4b**); data suggestive that EC2 might represent cells within the endothelial population that are actively undergoing remodeling.

**FIGURE 4:**
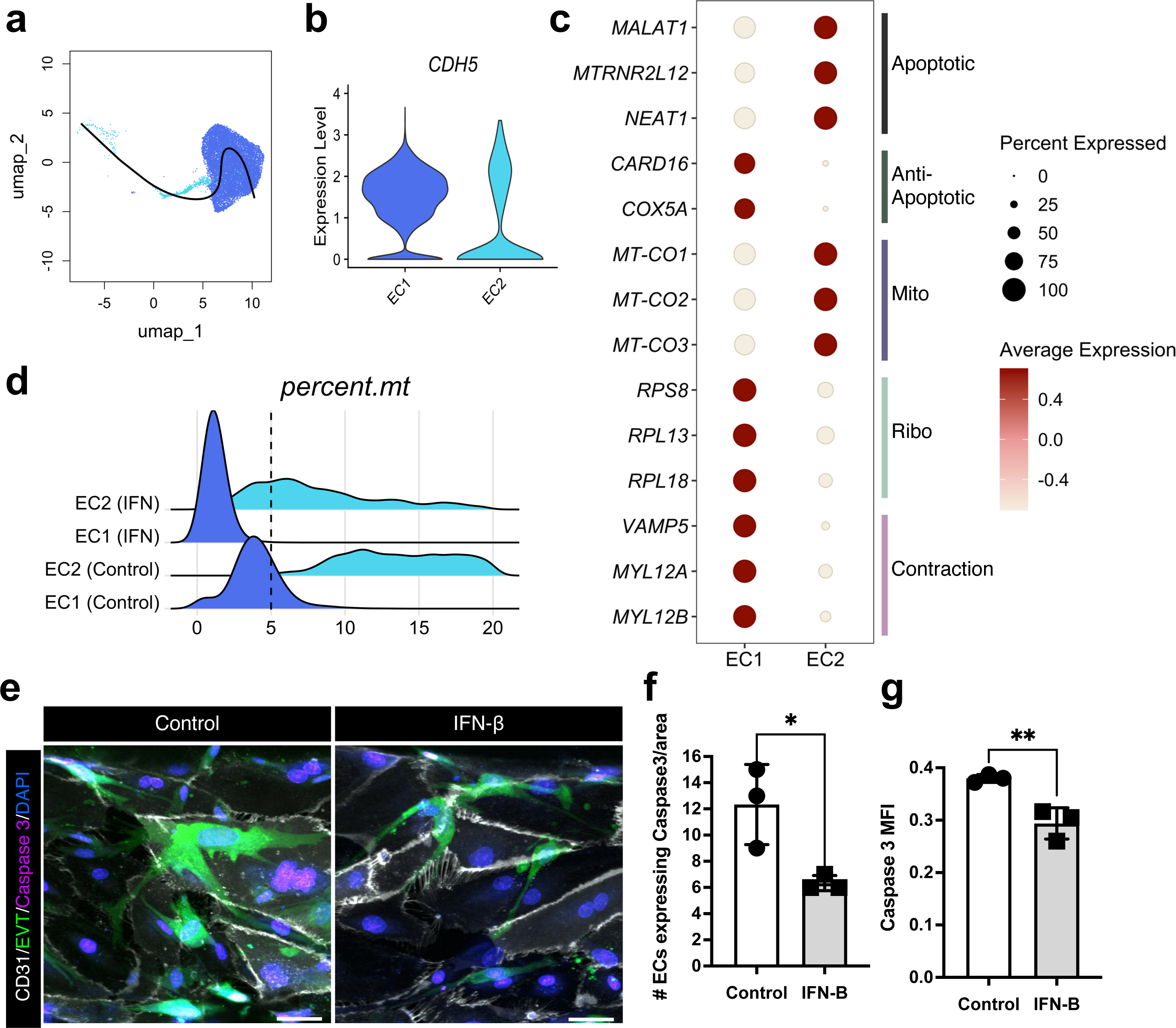
EVT-directed vascular remodeling is limited by type I interferon. **a.** Minimum spanning tree computed by Slingshot, visualized on the UMAP of EC subsets; EC1 (dark blue) and EC2 (light blue). **b.** Violin plots showing normalized expression of VE-Cadherin (CDH5) in individual EC subsets (x-axis). **c.** Dot plots showing normalized, log-transformed, and variance-scaled expression of genes characteristic of progression toward an apoptotic state and endothelial function (y-axis) in individual EC subsets (x-axis). **d.** Percent mitochondrial gene expression (x-axis) in individual EC subsets, separated by control (bottom) and IFN stimulated cells (top) (y-axis). **e.** Visualization and **f-g.** quantification of caspase-3 (magenta) expression. Scale bars, 50 μm. The representative images are from three independent experiments. Two-sided t-test (*n* = 3 independent devices per group). Data are presented as mean ± SEM. *p* values (. * = p < 0.05; ** = p < 0.01) are shown on graphs.

To explore further, we next evaluated differentially expressed genes in EC2s as compared with EC1s. Among the most upregulated genes in EC2s were various apoptosis regulatory long-coding RNAs such as *MALAT1, MTRNR2L12,* and *NEAT1* (**Figure 4c**). Consistently EC2s had decreased expression of anti-apoptotic genes, where we saw lower levels of the caspase inhibitor CARD16 (*CARD16)* and the anti-apoptotic (*COX5A*)^30,31^. In further support of EC2s transitioning to increased apoptotic state, we observed enhanced expression of mitochondrial cytochrome c subunits (*MT-CO1, MT-CO2, MT-CO3*)^32^ and reduced expression of various ribosomal proteins (*RPS8, RPL13, RPL18*)^33^. Lastly, we noted that EC2s had lower levels of genes associated with contraction (*VAMP5, MYL12A, MYL12B*) as compared with EC1s, indicating differences in EC functionality. Collectively, the gene expression signature of EC2s supports that this population is undergoing apoptotic remodeling.

We next assessed if IFN-β exposure ultimately impacted EVT-mediated apoptosis by evaluating the percent mitochondrial RNA in EC clusters within our sequencing data, a proxy for apoptotic cell fate^34^. We observed increased percentage of mitochondrial genes in both control EC clusters as compared to IFN-exposed EC clusters (**Figure 4d**), suggestive of less apoptotic remodeling in IFN-exposed ECs. Next, we aimed to corroborate this finding of distinct cell death between conditions by staining the IOC device for activated, cleaved caspase-3 (**Figure 4e**). Consistently, we found that IFN-β treatment limited EVT-mediated apoptosis within the IOC device which significantly reduced the amount of cleaved caspase-3 signal in the endothelial channel (**Figure 4f-g**). Cumulatively, these data indicate that this EVT-driven EC remodeling was limited when the IOC device was exposed to type I IFN exposure.

## DISCUSSION

Elevated type I IFN during embryo implantation is associated with abnormal placentation and adverse pregnancy outcomes, suggesting a role for type I IFN in affecting EVT function. We designed this study to evaluate the consequence of unwarranted systemic IFN on EVT function using a biomimetic model of human implantation in an organ-on-a-chip device^8^. We found that elevated type I IFN limited EVT invasion and endovascular EVT emergence. In evaluation of IFN exposed EVTs, we uncovered that sustained IFN signaling stunted EVT epithelial-to-mesenchymal progression with mesenchymal genes considerably reduced in all IFN stimulated EVTs. Prior literature has indicated IFN signaling is important for proper endometrial decidualization and spiral artery remodeling, yet the functional contribution of IFN signaling during implantation remained undefined^35,36^. Intriguingly, taken together with our findings, we speculate that IFN signaling might fine tune EMT transition in EVTs and thereby putatively facilitate directionality of invasion. Yet, further studies are needed to directly address this model.

Additionally, our work intended to investigate the impact of unwarranted IFN on implantation in order to mimic systemically elevated IFN in the environmental milieu, as seen in SLE patients. As such, one limitation to our study is that we only transcriptionally assessed one timepoint with a single, high dose of IFN, thereby limiting our ability to distinguish nuances of dose. Further, our current experimental set-up lacks the presence of IFN-responsive immune cells that have known functions in facilitating EVT invasion and maternal spiral artery remodeling, such as uterine NK cells^36,37^.

Although our findings describe a change in EVT trajectory and state after elevated IFN exposure, our results do not rule out a role for endothelial-specific IFN response on EVT invasion. Previous work with the IOC device indicated a critical role for intercellular communication between maternal endometrial EC in mediating directional EVT migration and invasion, likely through secretion of an unknown factor(s)^8^. Our studies did not address if IFN in ECs might impact this critical intercellular communication. Future work is needed to decipher individual contributions of cell types to the altered EVT function triggered by IFN signaling.

We identified a PE-like signature in IFN-EVTs compared with controls. PE is a life-threatening hypertensive disorder of pregnancy that confers a long-term, increased risk of cardiovascular disease in both mother and child^38^. PE is a multifactorial disease with multicomponent risk; thereby in order to effectively combat, there is a need to distinguish underlying mechanisms of the various risk factors. Currently, PE is understood to occur in two-stages with insufficient implantation and vascular remodeling leading to reduced placental perfusion (stage 1) that can lead to clinical manifestations of disease (stage 2)^39^. Interestingly, we found induction of genes that are implicated in both stages of PE. We found IFN stimulation induced expression of known PE risk factors, such as complement activation. Aberrant activation of the complement system during first trimester is associated with onset of PE later in gestation and thereby considered a stage 1 risk factor^27,28^. Furthermore, we identified noteworthy overlap with PE dysregulated genes that are known to be increased in stage 2 of PE, such as SAT1. SAT1 has been bioinformatically associated with PE physiopathology multiple times throughout literature, although the underlying mechanism remains poorly explored^40,41^. Overall, our study strikingly implicates unwarranted IFN signaling as a maternal disturbance that could trigger PE.

Glucocorticoids are steroid drugs with potent anti-inflammatory properties that function by targeting the type I and type II IFN pathways for immunosuppression. Multiple studies have attempted to utilize glucocorticoids to improve pregnancy outcomes in high-risk patients. Specifically, studies have suggested glucocorticoid treatment may be beneficial in specific patient groups, such as women with recurrent implantation failure after IVF^42,43^ and in women with idiopathic recurrent miscarriage^44^. Yet, there is little understanding of which patients may benefit from this intervention and the underlying mechanism(s) of action. Significantly, our data provides mechanistic insight to this gap in knowledge and specifically suggests temporal use of glucocorticoids during implantation and placentation may be beneficial for patients who present with elevated systemic IFN in order to combat pregnancy complications, such as development of PE.

Overall, our work implicates elevated systemic type I IFN as a maternal disturbance that can result in abnormal EVT function that could instigate PE.

## MATERIALS AND METHODS

### Obtaining human samples

Placental villi were obtained per protocol from first-trimester pregnancy terminations performed at the Penn Family and Pregnancy Loss Center (IRB#827072) (n=3, gestational ages 6-11 weeks). All subjects were counseled appropriately and provided written informed consent. Patients with preexisting medical conditions known to be associated with poor reproductive outcomes or any pregnancy complications in the current pregnancy were excluded from the study. Collected tissue was kept on ice and cell isolation was carried out within 1 hour of procurement.

### Primary cell isolation and culture

Primary Human Extravillous Trophoblasts: Primary EVTs were isolated from first trimester termination tissue based on an EVT-outgrowth based protocol established by Graham et al^45^. Briefly, villous tissue was finely minced and cultured at 37°C in RPMI 1640 medium containing 20% charcoal-stripped fetal bovine serum (FBS, Gibco, Catalog # 10437). After villous fragment attachment, EVT outgrowth occurred, and cells were separated from tissue during washing and passaging of the cells. The isolated EVTs were then cultured in 6 well plates using RPMI 1640 medium containing 20% FBS and 1% penicillin (100 U/ml)/streptomycin (100 U/ml). The cultured EVTs were used for constructing the implantation-on-a-chip model within the first 3 passages. EVT identity of the cells used in our experiments were characterized by immunostaining of cytokeratin-7 and HLA-G. Primary Human Endometrial Endothelial cells: Human endometrial microvascular endothelial cells were purchased (HEMEC, ScienCell, catalog #7010) and cultured in Endothelial Cell Growth Medium MV2 (EGM-MV2, Promocell, catalog # C-22022).

### Implantation-on-chip model

The 3-chamber device used to physiologically and spatially model EVT invasion is originally described and validated in a prior publication^37^. In brief, the fully enclosed device contains three parallel chambers, each individually accessible to fill with media and/or cells of choice. In our model, the middle chamber contains an extracellular matrix (ECM) to represent the endometrial stroma Primary human EVTs are in one side chamber (the “fetal” chamber), and human endometrial endothelial cells in the other (the “vascular” chamber), across the ECM. The arrangement allows in vivo-like directional migration of EVTs towards a micro-engineered maternal vessel to represent a physiologically relevant human model system (**Figure 1a-d**).

The ECM is a hydrogel precursor solution composed of a rat tail collagen Type-1 and Matrigel. Maternal endothelial cells and extravillous trophoblasts were cultured in their respective chambers, with media replenished every day. PromoCell Endothelial Cell Growth Medium MV2 (ready to use) media (C-22022) was used for the vascular chamber, and RPMI + 2 % FBS was used for the fetal chamber. The flow of media was prevented to simulate occluded blood flow at the maternal-fetal interface, with extra precautions taken during media exchange to minimize disruption of the culture environment. EVTs were fluorescently labelled with 5 μg/ml of CellTracker Green CMFDA (Thermo Fisher Scientific, Waltham, MA, USA, catalog # C7025) per manufacturer protocol in DMEM supplemented 2 % (v/v) FBS for 15 min at 37 °C, respectively. The device was incubated in a cell culture incubator in normoxic conditions maintained at 5% CO2 and 37 °C. After an initial 48 hours, the fetal and vascular chambers were treated with 100 or 1000 IU/mL of IFN-β with respective media and replaced every 24 hours (**Figure 1c**). Devices were imaged at 24, 72 and 120 hours after addition of IFN-β. After completion of cell culture experiments, media were removed from the reservoirs, and the channels were washed 3 times by flowing PBS. Cells cultured in flasks were similarly treated. Cells were fixed by introducing 4% paraformaldehyde (PFA, Thermo Scientific, catalog # AAJ19943K2) into the channels and incubated for 30 min at room temperature (RT), followed by 3 washes with PBS for all immunofluorescence experiments. After cell permeabilization and blocking with 0.1% Triton-X and 1% bovine serum albumin (BSA, Sigma, catalog # 5217), respectively, the cells were incubated with primary antibodies against Caspase-3 (Asp175, Cell Signaling, catalog # 9661), PECAM1 (2HB, Developmental Studies Hybridoma Bank) and Ki67 (37C7-12, Abcam, catalog # ab245113) diluted at 1:200 in 1% BSA-containing PBS solution at 4°C overnight. Subsequently, the cells were washed 3 times by flowing PBS and treated with appropriate secondary antibodies diluted at 1:200 in PBS containing 1% BSA for 2 hours at room temperature. The cells were counterstained with DAPI, then washed with PBS three times. Fluorescence imaging was performed using an inverted microscope with confocal capabilities (LSM 800, Carl Zeiss, Germany; 10X 0.45 NA objectives).

### Quantification of EVT invasion

Fluorescence images of EVTs were obtained from the ECM matrix region between the maternal vascular and fetal compartments of the implantation-on-a-chip device. High magnification images were collected from three separate devices per each experimental group with three representative images per device. EVT invasion was quantified at 24, 72 and 120 hours after addition of IFN-β, by (i) the number of invading EVTs and (ii) the area of EVT invasion. To evaluate the cell number, EVTs in the ECM compartments were manually counted, and the average of total cell counts was plotted. The invasion depth was also calculated at 120 hours, and quantified by the sum of the vertical travel distances of all EVTs in a given field of view was divided by the total number of EVTs in the same imaging area to calculate the average depth of invasion per cell number. Each data point in the invasion depth plots represents the mean of the average invasion depths measured from three randomly selected areas within a single device. Analysis of invasion area was achieved by using the Analyze pixels function of ImageJ (NIH) with appropriate thresholding to measure the area of ECM hydrogel covered by invading EVTs. A one-way ANOVA was performed to compare means, and a p-value <0.05 was considered statistically significant.

### Single cell RNA sequencing

Six IOC experiments per patient per condition (Control vs IFN-β) were run to analyze scRNA-sequencing, for a total of 36 devices. The IFN-β experiments used a dose of 1000 IU/mL of IFN-β based on the results seen in the invasion studies and increased the likelihood of establishing a difference between experimental groups. To assess the cells in the devices at the end of experiments, media was removed, and the devices were washed twice with PBS. After removing residual PBS, the reservoirs of each channel were filled with 0.25% Trypsin and incubated at 37 °C for 15 minutes while pipetting every 5 min until the ECM hydrogel in the middle channel was completely dissolved. Cell suspensions were collected, centrifuged at 300 × g for 5 min, and then the supernatant was removed using a pipette, leaving only the cell pellet. Cells were then washed in MACS buffer (sterile PBS, 1% FBS, 2 mM EDTA).

Next-generation sequencing libraries were prepared using the 10x Genomics Chromium Single Cell 3’ Reagent kit v3 as per manufacturer’s instructions. Libraries were uniquely indexed using the Chromium i7 Sample Index Kit, pooled, and sequenced on the Illumina NovaSeq6000 sequencer in a paired-end, single indexing run. Sequencing for each library targeted 20,000 mean reads per cell. Data was then processed using the Cellranger pipeline (10x genomics, v.6.0.0) for demultiplexing and alignment of sequencing reads to the GRCh38 transcriptome and creation of feature-barcode matrices.

Raw data was processed with the *DropUtils* R package^46,47^ to remove empty cells, and the *DoubletFinder* R package^48^ to remove probable doublets. Downstream analysis was done using *Seurat* (v.4.0.4) in R. Cells included in the analysis had greater than 500, and less than 15000 transcripts, greater than 250 and less than 5000 detected genes, a “complexity” score (log10GenesPerUMI) greater than 0.8, and a mitochondrial gene expression ratio of less than 20%. Cells were normalized for cell cycle phase and mitochondrial content using the *sctransform* function. Cells were integrated using a canonical correlation analysis to expect similar biological states among at least a subset of single cells across the conditions. Clustering was achieved using the *FindClusters()* function in Seurat. This constructed a K-nearest-neighbors graph of cells, using between 1 and 20 canonical correlation vectors, and partitioned them into ‘quasi-cliques’ based on similar gene expression. A clustering resolution of 0.2 was used to optimize the biologic variability expected based on prior studies based on a *clustertree* analysis. Differentially expressed genes (DEGs) were determined via the *FindMarkers()* in Seurat. DEGs were considered significant if they were increased or decreased by at least 1 log2 fold change, utilizing a false detection rate of 5%. *Monocle3* and *slingshot* R packages^49^ were used for pseudotime analysis of single-cell gene expression changes as a function of location and nearby cells.

For benchmarking purposes, we used sctransform and canonical correlation analysis in *Seurat* to further integrate scRNA-seq data of endovascular and interstitial EVTs from a study by Arutyunyan et al^12^. This reference data is of Donor P13, and publicly available via at https://www.reproductivecellatlas.org.

---

**Supplemental Figure 1: Additional characterization of implantation-on-a-chip (IOC) device.** Two days after seeding, we began daily treatments with a lower IFN-β (100 IU/mL) by supplementing the daily media exchange of the IOC device. We next assessed the properties of EVT invasion into the ECM toward the endothelial chamber for five days (**Figure 1c**). **a.** Quantification of EVT area invasion at Day 1, 3 and 5 after treatment with IFN-β (100 IU/mL). **b.** Quantification of depth of invasion after treatment with IFN-β (100 IU/mL). **c.** EVT proliferation was assessed by immunostaining of Ki-67 expression at 3- and 5-days post IFN-β (1000 IU/mL) exposure as compared with control and quantified. Data are presented as mean ± SEM. One-way ANOVA with Tukey’s multiple comparison test (*n* = 3 independent devices per group) (**a. & c.**) Two-sided t-test (*n* = 3 independent devices per group) (**b.**) **\*** = p < 0.05; **\*\*** = p < 0.01; **\*\*** = p < 0.001; **\*\*\*\*** = p < 0.0001 (*n* = 3 independent devices per group).

**Supplemental Figure 2: Integration analysis supports endovascular emergence in implantation-on-chip device.** To further benchmark EVT4s, we integrated snRNA-seq data from ∼2000 interstitial and endovascular EVTs from a reference human implantation site^12^. Visualized is the distribution of new clusters in our original clusters (**Figure 1**) and with reference data alone. Re-clustering with all cells in the UMAP confines over 90% the reference cells to cluster 3. In control cells, EVT4s are majority cluster 3. Consistent with our conclusions, we see loss of cluster 3 with IFN treatment, where cluster 3 in control cells outnumber cluster 3 IFN in cells by a factor of ∼2 proportionally and ∼3 in absolute counts (data not shown). Presented as bar plot representing the proportion (%) of EVT subsets separated by control and IFN stimulated cells.

**Supplemental Figure 2: Monocle3 analysis.** Minimum spanning tree computed by Monocle3, visualized on the Uniform manifold approximation and projection (UMAP) of EVT subsets; separated by control and type I IFN stimulated cells. In Monocle3, UMAP dimensionality reduction completed upstream of the trajectory inference that was computed by Monocle3. Consistent with our conclusions, Monocle3 predicted 4 clusters in control, but only 3 clusters in IFN thereby suggesting less heterogeneity upon IFN treatment.

## ACKNOWLEDGEMENTS

We thank the members of the Jurado laboratory for helpful discussions on this study. We acknowledge support of the Center for Applied Genomics core facility at the Children’s Hospital of Pennsylvania.

We gratefully acknowledge support from the Chan Zuckerberg Initiative (Single Cell Analysis of Inflammation; K.A. Jurado, M. Mainigi and D.D. Huh) in addition to the following agencies/foundations: Clinical and Translational Science Award TL1TR001880 to M.K. Simoni; University of Pennsylvania Presidential Fellowship to S.G. Negatu; HHMI Gilliam Fellowship to M.A. Arreguin and The Pew Charitable Trust (Biomedical Fellow) and Burroughs Wellcome Fund (Next Gen Pregnancy) to K.A. Jurado.

## AUTHOR CONTRIBUTIONS

M.K. Simoni: methodology, investigation, data curation, formal analysis, and writing-original draft. S.G. Negatu: data curation, formal analysis, visualization, and writing-review & editing. J.Y. Park: investigation, data curation, and formal analysis. S. Mani: methodology, investigation, and writing-review & editing. M.C. Arreguin: investigation and writing-review & editing. K. Amses: formal analysis. D.D. Huh: methodology, resources, supervision, and writing-review & editing. M. Mainigi: conceptualization, methodology, funding acquisition, resources, supervision, and writing-review & editing. K.A. Jurado: conceptualization, methodology, data curation, formal analysis, funding acquisition, resources, supervision, and writing-original draft.

## AUTHOR NOTES

Disclosures: The authors declare no competing interests exist.

## Notes

### Competing Interest Statement

The authors have declared no competing interest.

