## Supplemental Figures for "Type I interferon alters invasive extravillous trophoblast function"

### SUPPLEMENTAL FIGURE 1: Additional characterization of implantation-on-a-chip (IOC) device.

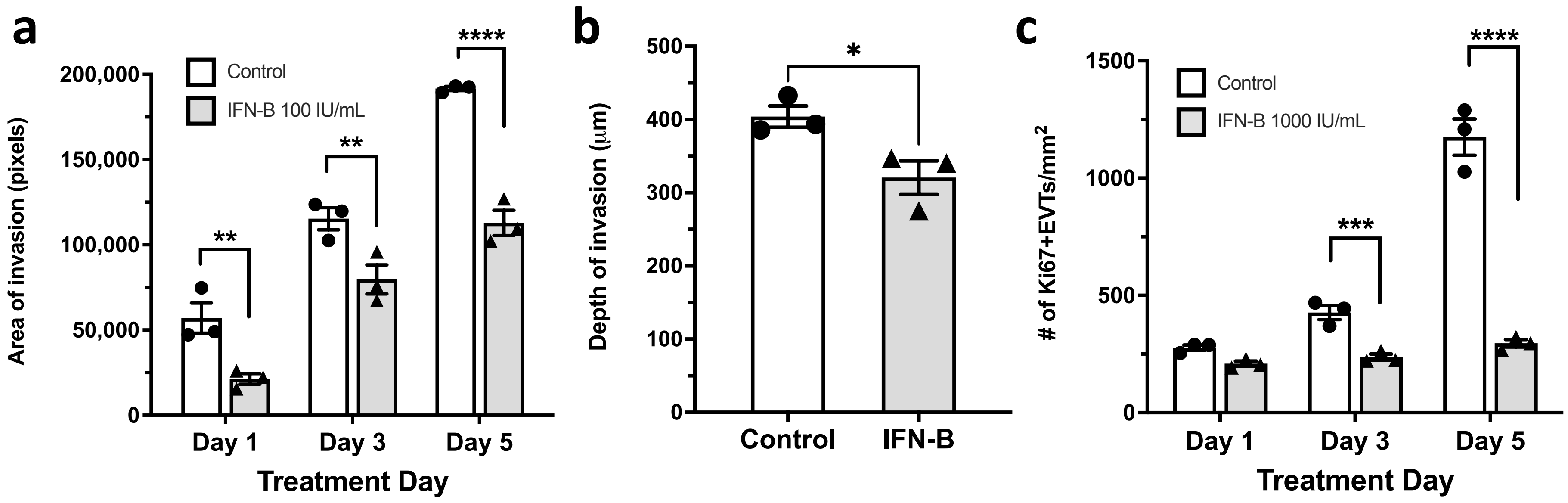

### SUPPLEMENTAL FIGURE 2: Integration analysis supports endovascular emergence in implantation-on-chip device.

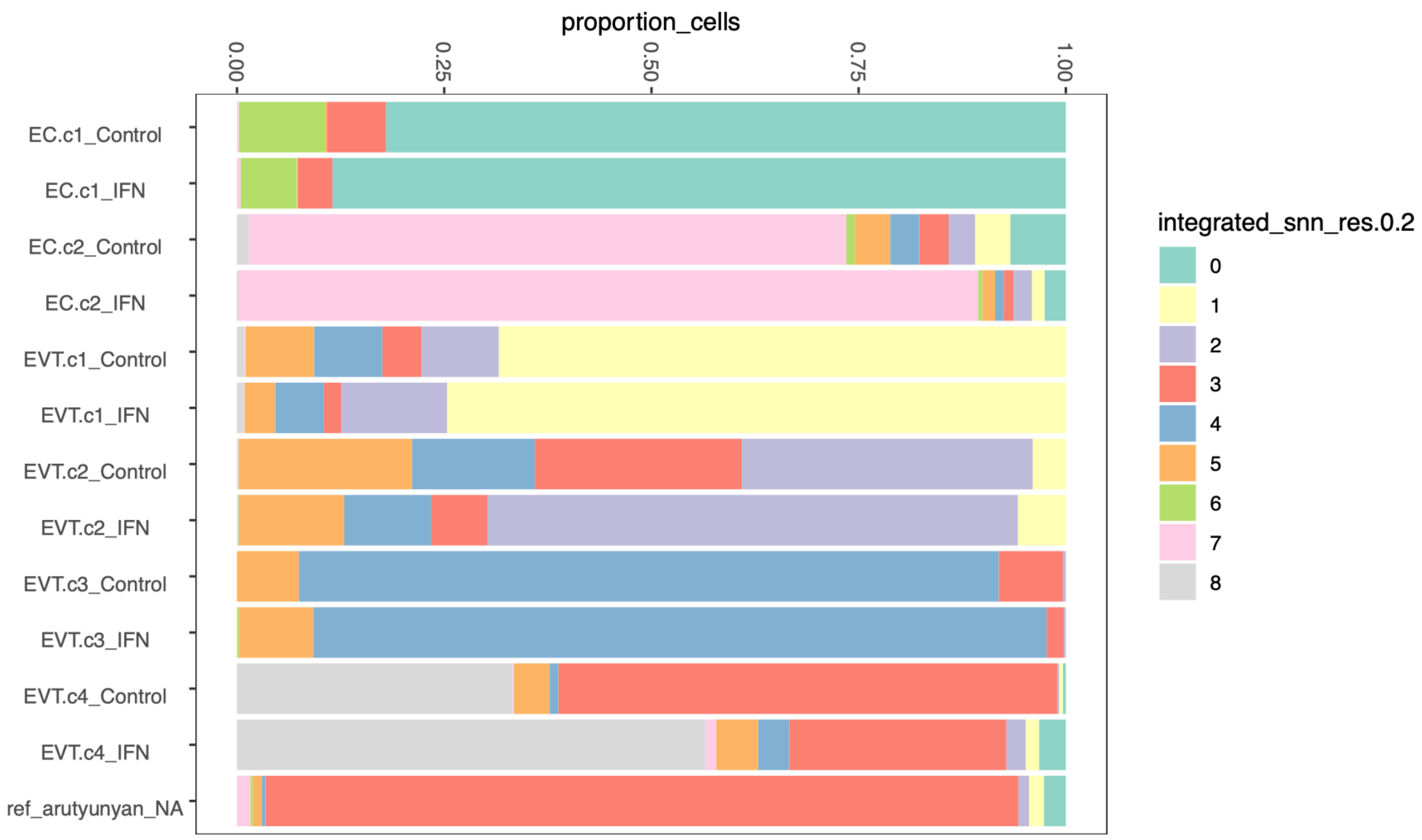

### SUPPLEMENTAL FIGURE 3: Monocle analysis

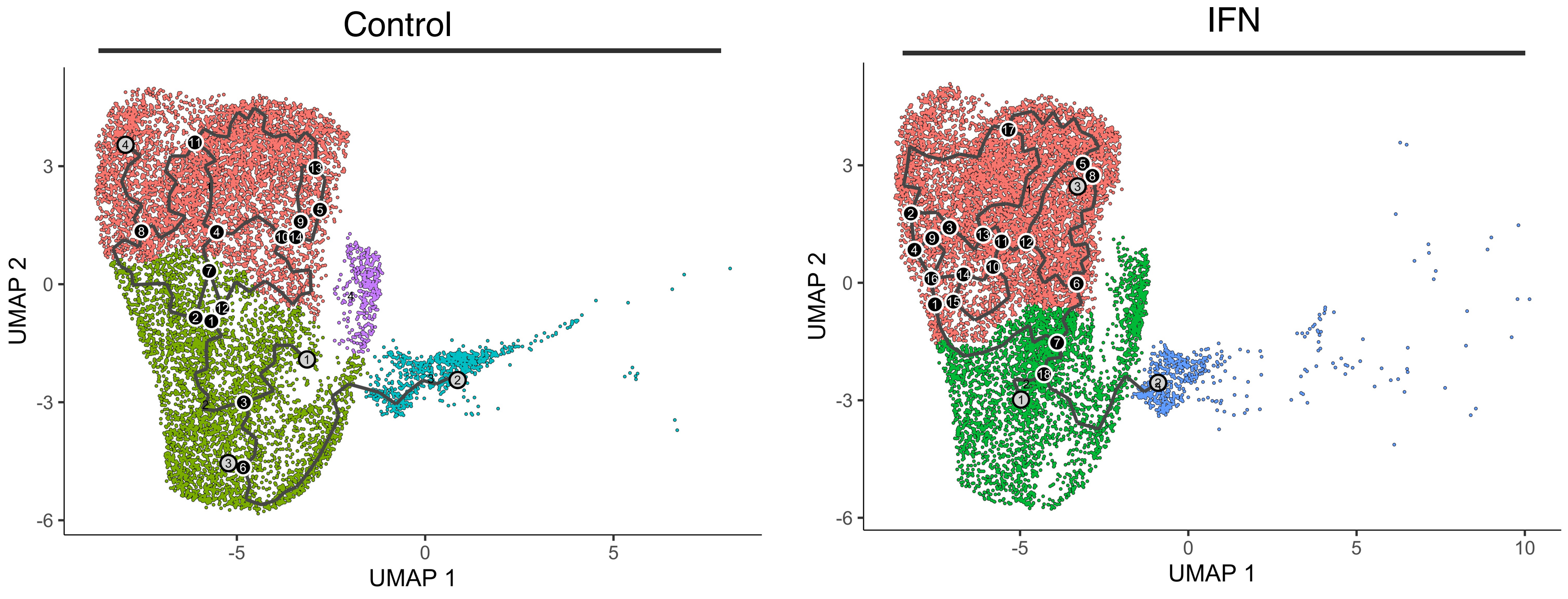
